# Proteasome dysfunction disrupts adipogenesis and induces inflammation via ATF3

**DOI:** 10.1101/2022.01.14.476367

**Authors:** Nienke Willemsen, Isabel Arigoni, Maja Studencka-Turski, Elke Krüger, Alexander Bartelt

## Abstract

**Objective:** Regulation of proteasomal activity is an essential component of cellular proteostasis and function. This is evident in patients with mutations in proteasome subunits and regulators, who suffer from proteasome-associated autoinflammatory syndromes (PRAAS). These patients display lipodystrophy and fevers, which may be partly related to adipocyte malfunction and abnormal thermogenesis in adipose tissue. However, the cell-intrinsic pathways that could underlie these symptoms are unclear. Here, we investigate the impact of two proteasome subunits implicated in PRAAS, Psmb4 and Psmb8, on differentiation, function and proteostasis of brown adipocytes.

**Methods:** In immortalized mouse brown pre-adipocytes, levels of *Psmb4, Psmb8*, and downstream effectors genes were downregulated through reverse transfection with siRNA. Adipocytes were differentiated and analyzed with various assays of adipogenesis, lipogenesis, lipolysis, inflammation, and respiration.

**Results:** Loss of Psmb4, but not Psmb8, disrupted proteostasis and adipogenesis. Proteasome function was reduced upon Psmb4 loss, but partly recovered by the activation of Nuclear factor, erythroid-2, like-1 (Nfe2l1). In addition, cells displayed higher levels of surrogate inflammation and stress markers, including Activating transcription factor-3 (Atf3). Simultaneous silencing of *Psmb4* and *Atf3* lowered inflammation and restored adipogenesis.

**Conclusions:** Our study shows that Psmb4 is required for adipocyte development and function in cultured adipocytes. These results imply that in humans with *PSMB4* mutations, PRAAS-associated lipodystrophy is partly caused by disturbed adipogenesis. While we uncover a role for Nfe2l1 in the maintenance of proteostasis under these conditions, Atf3 is a key effector of inflammation and blocking adipogenesis. In conclusion, our work highlights how proteasome dysfunction is sensed and mitigated by the integrated stress response in adipocytes with potential relevance for PRAAS patients and beyond.

**Highlights:**

- PRAAS-associated PSMB4 is required for brown adipocyte differentiation
- Loss of PSMB4 activates NFE2L1 to counteract proteasome dysfunction
- The ATF3 pathway regulates adipocyte dysfunction and inflammation
- Loss of ATF3 restores adipogenesis in cells with loss of PSMB4

**Graphical abstract:** 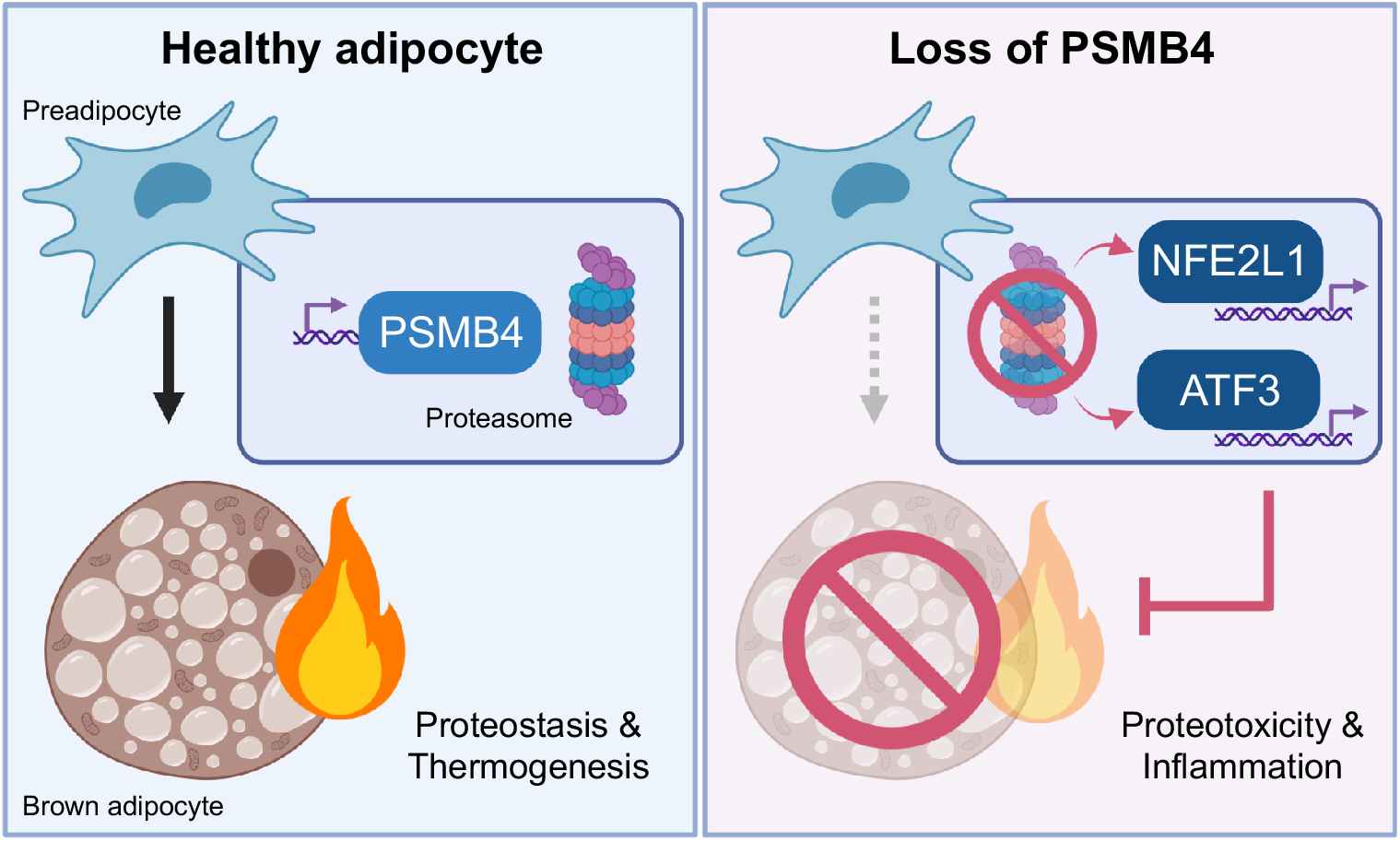

## 1. Introduction

Maintenance and regulation of the proteome by the ubiquitin-proteasome system (UPS) is essential for cellular function and health. Degradation of unwanted, obsolete, or damaged proteins by UPS is critical for cells to respond to environmental cues such as nutrients, temperature, or other forms of stress. Failure of the UPS is associated with severe consequences for the cells. In humans, this is evident in rare immunometabolic disorders caused by mutations in genes coding for proteasomal subunits or their assembly helper proteins. These diseases are characterized by a spectrum of rare auto-inflammatory syndromes known as proteasome associated auto-inflammatory syndromes (PRAAS), Chronic Atypical Neutrophilic Dermatosis with Lipodystrophy and Elevated Temperature (CANDLE) (1), Nakajo-Nishimura syndrome, or Joint Contractures-Muscular atrophy-microcytic anemia-Panniculitis-associated lipodystrophy (JMP) syndrome (2– 4). The first discovered cause of CANDLE/PRAAS was a missense mutation in the gene coding for proteasome subunit beta 8 (*PSMB8*, also known as *LMP7*) (1), and since then, other loss-of-function mutations have been revealed as underlying causes, including proteasome subunit beta 4 (*PSMB4*) (5). The disease symptoms in patients, which include sterile inflammation by increased production of type I interferons (IFNs) and lipodystrophy, are likely a complex result of disturbed UPS and maladaptive activation of the unfolded protein response (UPR) (6).

The CANDLE/PRAAS-related genes *PSMB4* and *PSMB8* are coding for proteasome subunits in the 20S core particle of the proteasome, which is a barrel-shaped structure of four heptameric rings of in total 28 subunits with a conserved modular architecture of α_1-7_β_1-7_β_1-7_α_1-7_. The active catalytic sites β1, β2, and β5 are located within this structure, which conceals them from the cytosol. The 20S core proteasome is associated with 19S regulator complexes to recognize, bind, and unfold ubiquitin-modified substrates for degradation. This allows for selective degradation of substrates that are translocated into the barrel (7). The modular structure of the proteasome is also evident in the diversity of isoforms with either alternative active sites or regulator complexes. The constitutive proteasome is ubiquitously found in most cell types and is responsible for the bulk of protein degradation, as well as controls many cellular pathways including signaling, metabolism, and proliferation (8). Immunoproteasomes bear alternative active sites, and are permanently expressed in cells of hematopoietic origin, but can be induced in response to interferons in other cell types (9). Immunoproteasomes are implicated in MHC-class I antigen processing, but also in many processes beyond the immune response, such as cell differentiation (9). Whereas PSMB4/β7, is a subunit of all proteasome isoforms, PSMB8/β5i, is an inducible subunit that replaces β5 in the constitutive proteasome when the proteasome is being remodeled to an immunoproteasome (10).

PRAAS patients display several pathologies with varying degrees of severity, yet aberrant inflammation and metabolic dysfunction are major hallmarks of the disease. Among these pathologies, the lipodystrophy phenotype is currently not well understood. Classically, lipodystrophy is caused by dysfunction of adipocytes, especially the ability to form healthy fat cells (11), but if PRAAS mutations cause cell-intrinsic differentiation defects or dysfunction of adipocytes has not been explored, yet. In addition, PRAAS patients display suffer from recurrent fever flares, which are thought to be caused by inflammation in the central nervous system. However, body temperature is a result of several mechanisms, including adaptive thermogenesis, and it is possible that aberrant thermogenesis is involved.

Brown adipose tissue (BAT) is a highly adaptive mammalian organ that is responsible for non-shivering thermogenesis (NST) (12). Stimulated by cold-induced norepinephrine (NE) and other stimuli, brown adipocytes generate heat through uncoupling protein-1 (Ucp1)and other thermogenic mechanisms. The herewith associated oxidative respiration is highly energy demanding, and is fueled by intracellular and circulating nutrients (13). Prolonged cold exposure leads to remodeling of BAT (12), which induces proteotoxic pressure on brown adipocytes (14). We have previously shown that an adaptive increase in proteasomal activity was essential for the maintenance of NST (15). Adapting proteasomal activity to the needs of the UPS is mediated by the transcription factor Nuclear factor erythroid-2, like-1 (Nfe2l1, also known as Nrf1 or TCF11). In most cell types, Nfe2l1 is continuously degraded by the proteasome, but it escapes its degradation when proteotoxic stress is increased in the cell, e.g., in the presence of chemical proteasome inhibitors. In that case, Nfe2l1 is cleaved from the ER membrane by the aspartyl protease DNA Damage Inducible 1 Homolog 2 (DDI2), translocates to the nucleus, and initiates the transcription of proteasome subunits, which results in restoration or heightening of proteasomal capacity (16,17). Lack of Nfe2l1 is associated with diminished NST, marked adipose tissue inflammation, and insulin resistance (15). Nfe2l1 was previously suggested to play a role in the disease mechanisms of CANDLE/PRAAS (6), but to the best of our knowledge no experimental evidence exists that Nfe2l1 is involved. Based on these findings, we hypothesized that compromised proteasome function, initiated by loss of CANDLE/PRAAS-related Psmb4 or Psmb8, impacts brown adipocyte proteostasis, development, function, and inflammation.

## 2. Materials and methods

### 2.1 Mice husbandry and tissue collection

All animal experiments were performed with approval of the local authorities (License: ROB-55.2-2532.Vet_02-30-32). Psmb8 whole-body knock-out mice were described previously (18,19). Animals were housed in individually ventilated cages at room temperature (22 °C), with a 12-h light–dark cycle, and fed chow-diet (Sniff) and water ad libitum. For cold exposure, wild-type C57BL/6J mice (Janvier) were housed at 30 °C or 4 °C for seven days. Mice were killed with cervical dislocation. Tissues were snap-frozen in liquid nitrogen and stored at −80 °C.

### 2.2 Cell culture, treatments, and reverse transfection

For cellular experiments, we used immortalized WT-1 mouse brown preadipocytes. WT-1 cells were maintained in DMEM Glutamax (Thermo Fisher, supplemented with 10 % v/v FBS (Sigma) and 1 % v/v PenStrep (Thermo), and incubated 37 °C, 5 % CO_2_. They were continuously kept between 20-80 % confluency and split two or three times per week. Cells were used until their passage number exceeded 18. For differentiation, WT-1 cells were grown to 90 % confluency. Then, they were differentiated in WT-1 induction medium (850 nM insulin (Sigma), 1 μM dexamethasone (Sigma), 1 μM T3 (Sigma), 1 μM rosiglitazone (Cayman), 500 nM IBMX (Sigma) and 125 nM indomethacin (Sigma)) from day 0 to day 2, and in WT-1 differentiation medium (1 μM T3 and 1 μM rosiglitazone) from day 2 up until day 5, with the medium being replaced every other day. For in vitro treatments, differentiated cells were treated with DMSO, 100 nM Epoxomicin (Millipore) for 6 h, 100 nM ONX0914 (Adooq) for 6 h, 1 μM NE (Sigma) for 1 h, or 1 μM CL316,143 (Tocris) for 16 h. For in vitro gene silencing, mRNA levels of target genes were knocked down through reverse transfection with SMARTpool silencing RNA (siRNA, Dharmacon) in Lipofectamine RNAiMAX transfection reagent (Thermo Fisher), used according to manufacturer’s instructions. siRNAs were added to the cells in 30 nM for single knock-down or two times 30 nM for double knockdown. Reverse transfection took place 1 day before induction. The transfection mix was replaced for standard induction medium 24 h after transfection. Cells were harvested as pre-adipocytes, early adipocytes, or mature adipocytes; on day 0, day 3 or day 5 of differentiation, respectively. If not further specified, assays were performed on mature adipocytes, i.e., day 5.

### 2.3 Gene expression analysis

We used NucleoSpin RNA kit (Macherey Nagel), according to the manufacturer’s instructions, for RNA extraction. RNA concentrations were then measured with a NanoDrop spectrophotometer (Thermo Fisher). Complementary DNA (cDNA) was prepared by adding 2 μl Maxima H Master Mix (Thermo Fisher) to 500 ng RNA, adjusted with H_2_O to 10 μl total. This cDNA mixture was diluted 1:40 in H_2_O. Relative gene expression was measured with qPCR. Per reaction, 4 μL cDNA, 5 μL PowerUp™ SYBR Green Master Mix (Applied Biosystems) and 1 μL of 5 μM primer stock (Sigma, see Supplementary table 1 for sequences) were mixed. Expression was measured in a Quant-Studio 5 RealTime PCR system (Thermo Fisher, 2 min 50 °C, 10 min 95 °C, 40 cycles of 15 s 95 °C, 1 min 60 °C). Cycle thresholds (Cts) of genes of interest were normalized to *TATA-box binding protein (Tbp)* levels by the ΔΔCt-method. Relative gene expression is displayed as fold change to the appropriate experimental control groups.

### 2.4 Protein isolation and analysis

The samples were lysed in RIPA buffer (50 mM Tris (Merck, pH = 8), 150 mM NaCl (Merck), 5 mM EDTA (Merck), 0.1 % w/v SDS (Carl Roth), 1 % w/v IGEPAL® CA-630 (Sigma-Aldrich), 0.5 % w/v sodium deoxycholate (Sigma-Aldrich)) freshly supplemented with protease inhibitors (Sigma-Aldrich) in a 1:100 ratio. Cell lysates were centrifuged for 30 min (4 °C, 21,000 *g*) and tissue lysates were centrifuged 3 times 30 min, before supernatant was collected. Protein concentrations were determined using the Pierce BCA assay (Thermo Fisher) according to the manufacturer’s instructions. 15-30 μg protein per sample were denatured with 5 % vol/vol 2-mercaptoethanol (Sigma) for 5 min at 95 °C before they were loaded in Bolt™ 4–12 % Bis-Tris gels (Thermo Fisher). After separation, proteins were transferred onto a 0.2 μm PVDF membrane (Bio-Rad) using the Trans-Blot® Turbo™ system (Bio-Rad) at 12 V, 1.4 A for 16 min. The membrane was briefly stained with Ponceau S (Sigma) to verify protein transfer and consequently blocked in Roti-Block (Roth) for 1 h at room temperature. The membranes were incubated overnight in primary antibody dilutions (1:1000 in Roti-block) at 4 °C. The primary antibodies used were: Beta-tubulin (Cell Signaling, 2146), Psmb4 (Santa Cruz, sc-390878), Psmb8 (Cell Signaling, 13635), Nfe2l1 (Cell Signaling, 8052), Ubiquitin/P4D1 (Cell Signaling, 3936), Ucp1 (Abcam, ab10983), and Proteasome 20S alpha 1+2+3+5+6+7 (Abcam, ab22674). After washing with TBS-T (200 mM Tris (Merck), 1.36 mM NaCl (Merck), 0.1 % v/v Tween 20 (Sigma)), the membranes were incubated in secondary antibody (Santa Cruz) solutions (1:10,000 in Roti-block) for 90 min at room temperature. The membranes were washed in TBS-T and developed using SuperSignal West Pico PLUS Chemiluminescent Substrate (Thermo Fisher) in a Chemidoc imager (Bio-Rad). We normalized the protein bands to β-tubulin with Image Lab software (Bio-Rad). The uncropped images are in Supplementary fig. S1.

### 2.5 Oil Red O staining

We used Oil Red O (ORO) staining to measure lipid content in adipocytes. Cells were washed with cold DPBS (Gibco), fixed in zinc formalin solution (Merck) for 60 min at room temperature, and then again washed the cells with DPBS. After the samples had completely dried, they were stained with ORO mix (60 % v/v Oil-Red-O solution (Sigma), 40 % v/v H_2_O) for 60 min. After the incubation time, the cells were washed several times with water. We took pictures to visualize lipid content. To quantify lipid content, the ORO staining was eluted with 100 % isopropanol, and the absorption was measured at 500 nm in a Tecan plate reader.

### 2.6 Free glycerol assay

To assess lipolysis, we used Free Glycerol Reagent (Sigma F6428) and Glycerol standard solution (Sigma G7793) to measure free glycerol concentrations in the cell culture supernatant. After treatment and just before harvesting the samples, we collected cell culture medium to measure free glycerol content. The kit was used according to manufacturer’s instructions.

### 2.7 Proteasome activity

To prepare lysates for the proteasomal activity assay, cells were lysed in lysis buffer (40 mM TRIS pH 7.2 (Merck), 50 nM NaCl (Merck), 5 mM MgCl_2_(hexahydrate) (Merck), 10 % v/v glycerol (Sigma), 2 mM ATP (Sigma), 2 mM 2-mercaptoethanol (Sigma). Activity was measured using the Proteasome Activity Fluorometric Assay II kit (UBPBio, J41110), according to manufacturer’s instructions in a Tecan Plate reader. This assay allowed for measurements of chymotrypsin-like, trypsin-like, and caspase-like activity. The results were then normalized to either protein, using Bio-Rad Protein Assay Kit II (Bio-Rad), or DNA, using the Quant-iT PicoGreen dsDNA assay kit (Invitrogen, P7589), both according to manufacturer’s instructions.

### 2.8 Native PAGE

The protocol for in-gel proteasome activity assay and subsequent immunoblotting was previously described in detail (20). In short, cells were lysed in Lysis buffer (50 mM Tris/HCl pH 7.5, 2 mM DTT, 5 mM MgCl_2_, 10 % glycerol (vol/vol), 2 mM ATP, 0.05 % Digitonin (v/v)), with a phosphate inhibitor (PhosphoStop, Roche Diagnostics). The suspensions were kept on ice for 20 min and then centrifuged twice. Protein concentration was determined with Bio-Rad Protein Assay Kit II and 15 μg sample protein was loaded in a NuPAGE 3-8 % Tris-Acetate gel (Thermo Fisher). The gel was run at a constant voltage of 150 V for 4 h. Afterwards, the gel was incubated in an activity buffer (50 mM Tris, 1 mM MgCl_2_, 1 mM DTT) with 0.05 mM chymotrypsin-like substrate Suc-Leu-Leu-Val-Tyr-AMC (Bachem) for 30 min at 37 °C. The fluorescent signal was measured using ChemiDoc MP (Bio-Rad). The gel was then incubated in a solubilization buffer (2 % SDS (w/v), 66 mM Na_2_CO_3_, 1.5% 2-Mercaptoethanol (v/v)) for 15 min to prepare the samples for blotting. Through tank transfer, the samples were transferred to a PVDF membrane by 40 mA, overnight and developed as described above. The uncropped images are in Supplementary fig. S1.

### 2.9 Extracellular flux analysis (Seahorse)

Mitochondrial respiration was measured with Seahorse Cell Mito Stress Test (Agilent) with some adjustments to the manufacturer’s protocol. Briefly, WT-1 cells were cultured in a 24-well Seahorse plate until day 3 of differentiation. Culture medium was then replaced for Seahorse medium (XF DMEM pH 7.4, 10 mM glucose, 1 mM Pyruvate, 2 mM L-glutamine) and the cells were incubated for 60 min more at 37 °C without CO_2_ before being placed in the Seahorse Analyzer XFe24. In the assay, the cells were treated with NE (final concentration in the well was 1 μM), oligomycin (1 μM), FCCP (4 μM) and μM rotenone–antimycin A (0.5 μM). The reagents were mixed for 3 min, followed by 3 min of incubation, and 3 min of measurements. Afterwards, total DNA was measured for normalization, with CyQUANT Cell Proliferation Assay Kit (Thermo Fisher), according to manufacturer’s instructions.

### 2.10 Statistics

Data were analyzed with ImageLab, Microsoft Excel and Graphpad Prism. Unless otherwise specified, data are shown as mean ± standard error of the mean (SEM), including individual measurements. Multiple student’s t-test was used for experiments when comparing two groups and one variable, one-way ANOVA with Bonferroni post-hoc test was used when comparing three or more groups, and two-ANOVA with Bonferroni post-hoc test was used for comparing two groups with two variables. P-values lower than 0.05 were considered significant and are as such indicated in the graphs with an asterisk or as different letters between groups.

## 3. Results

### 3.1 Psmb4 is induced during adipocyte differentiation and regulated by cold

Proteasome function is an evolutionary conserved feature of every mammalian cell, but little is known about the relative presence of proteasome subtypes and subunits throughout the body. We focused on a mouse brown adipocyte model, as this allows studying both differentiation and thermogenic function. To understand better if CANDLE/PRAAS-linked subunits are expressed in adipose tissue, we analyzed the gene expression of *Psmb4* and *Psmb8* and found that both are robustly expressed in BAT of mice housed at 30 °C thermoneutrality (Fig. 1A). As the expression of many genes in BAT is linked to activation status, for example cold-induced NST, we also determined the expression in BAT from cold-adapted animals. While *Psmb4* remained unchanged, *Psmb8* expression was lower at 4 °C compared to 30 °C (Fig. 1A). Next, we analyzed cultured brown adipocytes differentiated from immortalized preadipocytes. *Psmb4* mRNA levels were markedly higher in differentiated adipocytes compared to pre-adipocytes (Fig. 1B). In contrast, *Psmb8* expression was lower in differentiated adipocytes compared to pre-adipocytes (Fig. 1B). We also tested the response of these genes to pharmacological activation. In line with the in vivo data, *Psmb4* remained unchanged and *Psmb8* was lower in cells activated with NE or the β3-adrenergic agonist CL316,243 (Fig. 1C). In summary, these data show that both *Psmb4* and *Psmb8* are robustly expressed in adipocytes, but while *Psmb4* seems to be implicated in mature brown adipocyte function, *Psmb8* expression was diminished under these conditions. However, to rule out that Psmb8 might nevertheless have an important function we performed loss-of-function experiments. First, we collected adipose tissues from mice with whole-body deletion of Psmb8 (19) and analyzed gene expression and histology. Mice deficient in Psmb8 had no apparent BAT abnormalities, which also was reflected in unchanged gene expression (Fig. S2A). The notion that Psmb8 is dispensable was supported by in vitro experiments, as silencing of *Psmb8* neither in immortalized pre-adipocytes nor in mature adipocytes produced an impact on adipogenesis or stress markers (Fig. S2B-D). Also, there was no difference in lipid content in mature adipocytes (Fig. S2E). In conclusion, loss of Psmb8 did not affect adipogenesis or BAT phenotype in vitro or in vivo. The concomitant down-regulation of Psmb8 in response to β3-adrenergic stimulation may indicate that immunoproteasome function is dispensable in brown adipocytes.

**Fig. 1:**
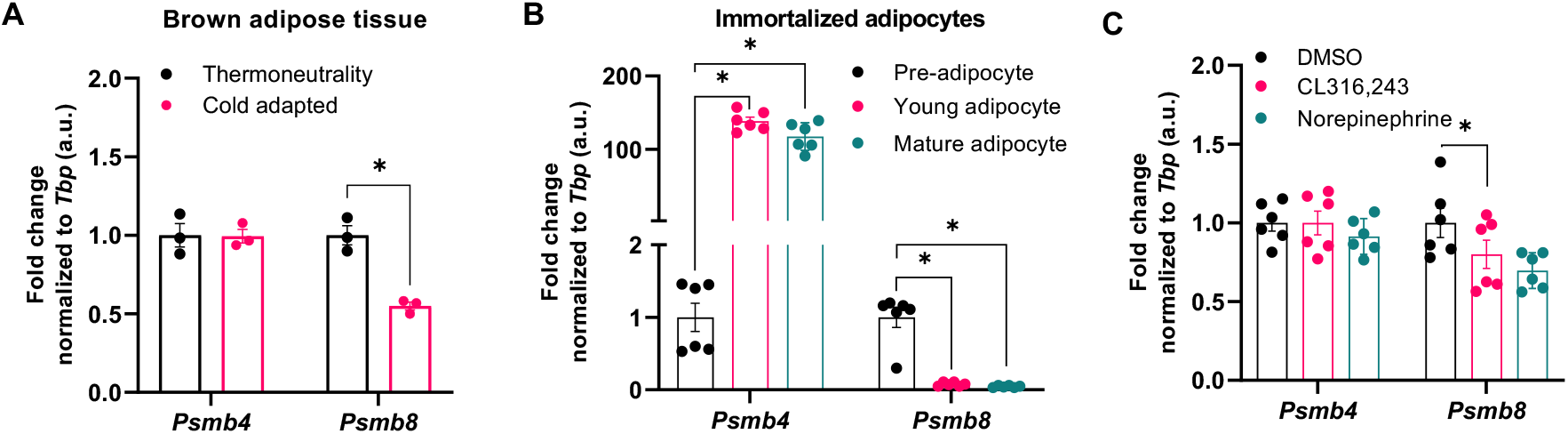
Regulation of *Psmb4* and *Psmb8* expression in brown adipocytes. **(A)** Relative gene expression in BAT from mice housed at thermoneutrality (30 °C) or cold (4 °C) for 7 d (n = 3 biological replicates). **(B)** Relative gene expression in immortalized WT-1 brown adipocytes at day 0, day 3 (young) or day 5 (mature) of differentiation, or **(C)** in adipocytes (day 5) after treatment with DMSO, 1 μM CL316,243 (16 h) or 1 μM NE (1 h). Unless otherwise specified: n = 6 independent measurements from 2 separate experiments. Data are mean ± SEM. Significant if *P* < 0.05, indicated by (*) or different letters.

### 3.2 Loss of Psmb4 disrupts adipogenesis and proteostasis

As Psmb4 was induced during adipocyte differentiation and highly expressed in activated brown adipocytes and BAT, we next studied the role of Psmb4 in brown adipocyte proteostasis, differentiation, and thermogenic function in more detail. Loss of Psmb4 was achieved by silencing *Psmb4* expression through siRNA (“siPsmb4 cells”). Transfection one day before the start of differentiation successfully reduced mRNA levels and this effect remained stable during different stages of adipogenesis (Fig. 2A), resulting in lower protein levels (Fig. 2B). To determine proteasome function in these cells we performed native PAGE analysis which allows examining activity and protein levels of macromolecular proteasome configurations, i.e., 30S, 26S, and 20S. Chymotrypsin-like peptide hydrolysis activity of the proteasome was lower in all three proteasome complexes in siPsmb4 compared to control cells (Fig. 2C). There were less 30S and 26S complexes present in siPsmb4 cells, as well as an increase in various subcomplexes, including precursors of the 20S proteasome (Fig. 2C). However, the overall levels of proteasome subunits were unchanged (Fig. 2D), indicating an assembly deficit of the proteasome in the absence of Psmb4. Interestingly, in cell lysates of siPsmb4 compared to control cells, proteasomal activity was lower in immature adipocytes but higher in mature adipocytes (Fig. 2F). However, in siPsmb4 cells overall ubiquitin levels were higher compared to control cells (Fig. 2E), which indicates a shift in the overall function of UPS. Interestingly, these changes in UPS were accompanied by higher levels of active Nfe2l1 in the siPsmb4 compared to control cells (Fig. 2F), indicating that the brown adipocyte mount an adaptive response to overcome UPS dysfunction. Regardless of this Nfe2l1 activation, silencing of Psmb4 led to lower levels of adipogenesis and thermogenesis markers during the early phase of adipocyte differentiation (Fig. 2G). In mature adipocytes, these changes were less pronounced albeit detectable for *Pparg* and *Ucp1* (Fig. 2H). Instead, *Ccl2*, a surrogate marker for the adipocyte stress response (21), was markedly higher in siPsmb4 compared to control cells (Fig. 2H). These phenotypic alterations translated into lower lipolysis after treatment CL316,243 (Fig. 2I). In summary, loss of Psmb4 led to proteotoxic stress through proteasome dysfunction as well as limited adipogenesis and adipocyte function.

**Fig. 2:**
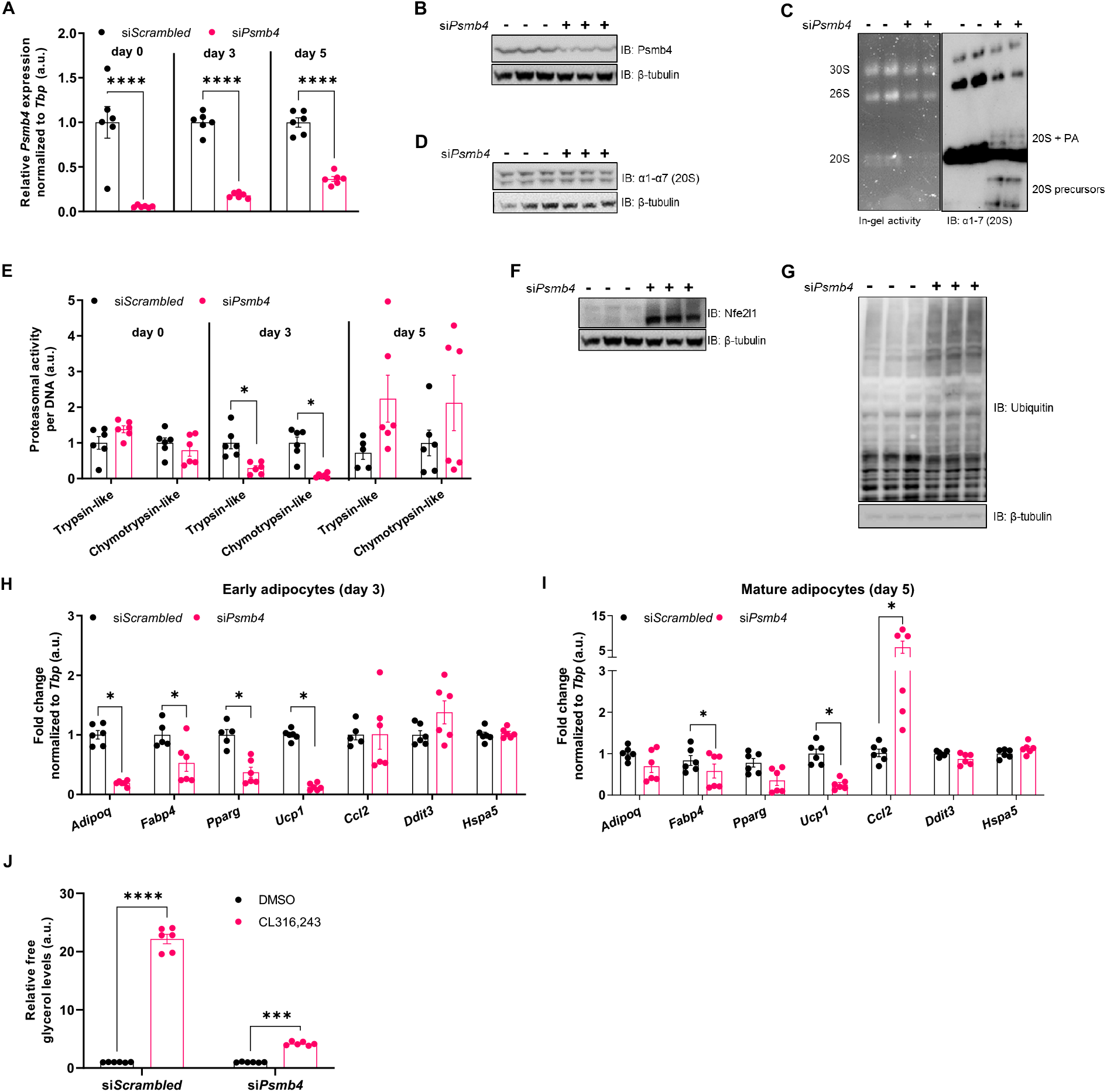
Psmb4 controls adipogenesis, adipocyte health and proteostasis. **(A)** Relative gene expression of *Psmb4* at different timepoints after knockdown with scrambled or *Psmb4* siRNA. **(B)** Representative immunoblot of Psmb4 in siScrambled and siPsmb4 adipocytes. **(C)** Native PAGE with in-gel chymotrypsin-like proteasome activity and immunoblot of α1-α7 (20S) proteasome subunits. PA = proteasome activators. **(D)** Immunoblot of α1-α7 (20S) proteasome subunits. **(E)** Trypsin-like and chymotrypsin-like proteasome activity in adipocytes (day 5) at different timepoints at different timepoints, normalized to DNA content. **(F)** Immunoblot of Nfe2l1. **(G)** Immunoblot of ubiquitin. **(H**,**I)** Relative gene expression of adipogenesis and stress markers adipocytes after *Psmb4* knockdown measured on day 3 (H) and day 5 (I) of differentiation. **(J)** Relative free glycerol response in adipocytes (day 5) after DMSO or 1 μM CL316,243 treatment for 3 h. Unless otherwise specified: n = 6 independent measurements from 2 separate experiments. Data are mean ± SEM. Significant if *P* < 0.05, indicated by (*) or different letters.

### 3.3 Activation of Nfe2l1 partially compensates for the loss of Psmb4

While loss of Psmb4 had a marked effect on proteasome function, it did not completely disable UPS. We hypothesized that the activation of Nfe2l1 mitigated some of the effects of Psmb4 silencing. Therefore, we tested whether the adipocyte phenotype was more severe in a double *Psmb4* and *Nfe2l1* knockdown model. Double knockdown successfully led to lower mRNA as well as protein levels of both Psmb4 and Nfe2l1 (Fig. 3A,B). Indeed, double knockdown of both Nfe2l1 and *Psmb4* knockdown led to lower proteasomal activity compared to control cells, or single silencing. (Fig. 3C). The effects of the double knockdown of *Psmb4* and *Nfe2l1* on adipogenesis markers compared to control or the single knockdown cells were minimal (Fig. 3D). However, *Psmb4* knockdown was associated with higher adipocyte inflammation surrogate markers compared to control cells, but this effect was not further amplified by additional *Nfe2l1* knockdown (Fig. 3E). In addition, loss of Psmb4 led to markedly higher mRNA levels of *Activating Transcription Factor-3 (Atf3)*, an important transcription factor linked to the integrated stress response, but not of other stress markers (Fig. 3D,F), an effect also seen in CANDLE/PRAAS patients (5). These changes were independent of *Nfe2l1* knockdown (Fig. 3D,F). In summary, activation of Nfe2l1 helps to sustain proteasome function upon loss of Psmb4, but this effect did neither restore adipogenesis nor reduce the adipocyte stress response under these conditions.

**Fig. 3.**
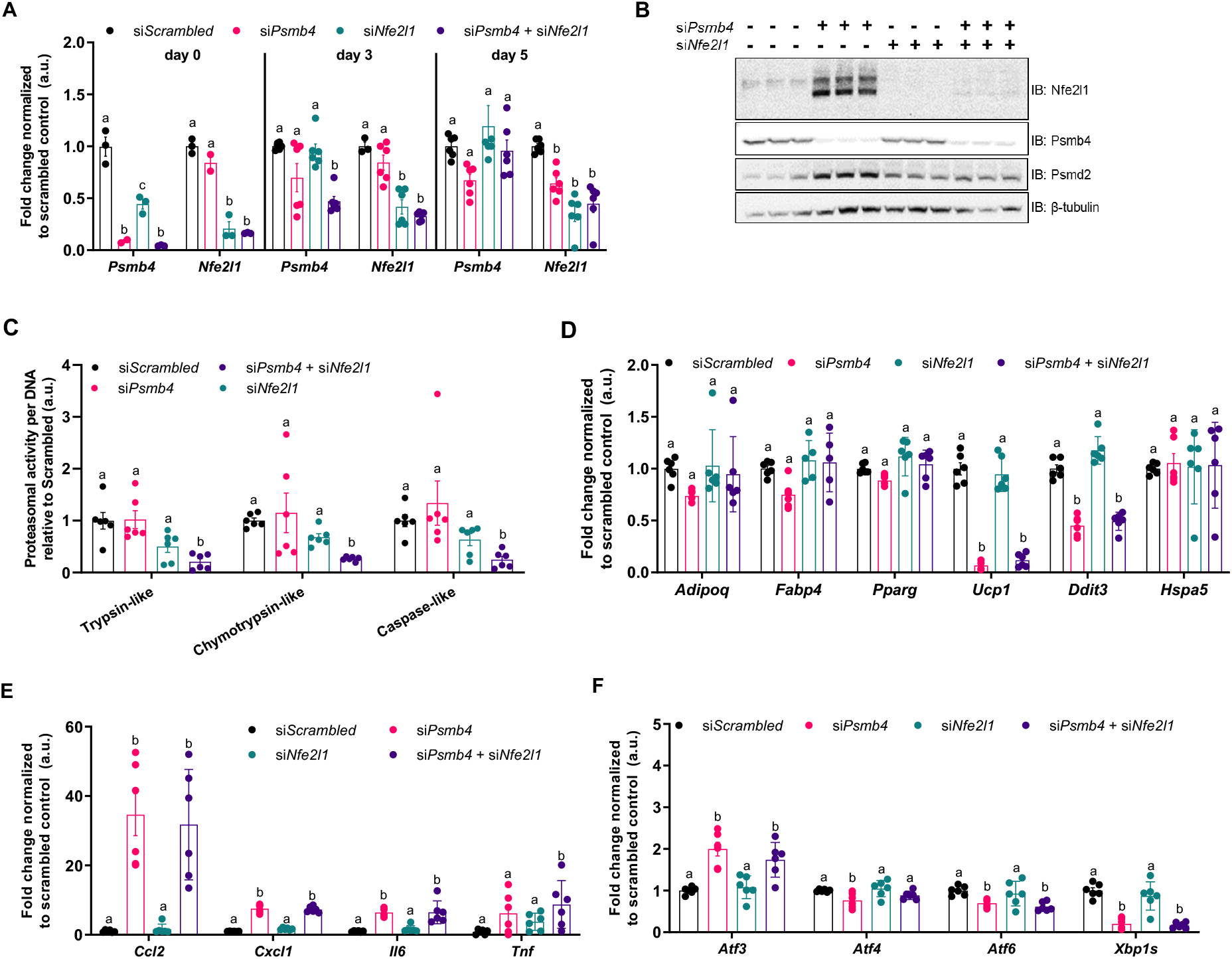
Loss of Psmb4 initiates a proteostatic stress-response via Nfe2l1. **(A)** Relative gene expression of *Psmb4* and *Nfe2l1* at different timepoints after knockdown with scrambled, *Psmb4* and/or *Nfe2l1* siRNA. **(B)** Representative immunoblot of Nfe2l1, Psmb4, Psmd2 and β-tubulin in siScrambled, siPsmb4, siNfe2l1 or siPsmb4 + siNfe2l1 adipocytes. **(C)** Trypsin-like, chymotrypsin-like, and caspase-like proteasome activity in adipocytes (day 5), normalized to DNA content. **(D)** Relative gene expression of adipogenesis and stress markers in adipocytes (day 5). **(E)** Relative gene expression of inflammation markers in adipocytes (day 5). **(F)** Relative gene expression of ER stress and unfolded protein response markers in adipocytes (day 5). Unless otherwise specified: n = 6 independent measurements from 2 separate experiments. Data are mean ± SEM. Significant if *P* < 0.05, indicated by (*) or different letters.

### 3.4 Atf3 links loss of Psmb4 to inflammation and adipogenesis

The fact that Atf3 expression was selectively induced in siPsmb4 cells caught our attention, as Atf3 is a stress-induced transcription factor that reportedly has been associated with adipogenesis and lipogenesis (22). To investigate the role of Atf3 in our model system, we silenced *Psmb4, Nfe2l1* or *Atf3* individually and in a paired fashion (Fig. 4A). This approach did not affect the viability of the adipocytes (Fig. 4B). Unlike targeting Nfe2l1 (Fig. 3C), *Atf3* knockdown in addition to *Psmb4* knockdown did not affect proteasomal activity (Fig. 4C). However, *Atf3* knockdown in addition to *Psmb4* knockdown normalized inflammation markers compared to siPsmb4 and control cells (Fig. 4D). In line with this normalized gene expression pattern, double knockdown of both *Psmb4* and *Atf3* largely rescued lipogenesis and restored lipolysis as follows. Particularly, cells with double knockdown of *Psmb4* and *Atf3* displayed markedly higher lipid content compared to siPsmb4 cells, and similar lipid content compared to control cells (Fig. 4E). Knockdown of only *Atf3* or *Nfe2l1* had no effect. Brown adipocytes with double knockdown of *Psmb4* and *Atf3* displayed higher NE-induced glycerol release compared to siPsmb4 cells, almost as much as control cells (Fig. 4F). Again, knockdown of only *Atf3* or *Nfe2l1* had no effect. (Fig. 4F). Finally, to assess NST, we measured NE-stimulated oxygen consumption followed by a mitochondrial stress test using a Seahorse Analyzer. In line, the finding that loss of Psmb4 diminishes brown adipocyte function, siPsmb4 cells displayed abolished NE-stimulated and uncoupled respiration as well as lower maximal capacity compared to control cells. Double knockdown of both *Psmb4* and *Atf3* largely rescued these alterations in mitochondrial function (Fig. 4G). Knockdown of only *Atf3* or *Nfe2l1* had no effect (Fig. 4G). In conclusion, loss of Psmb4 disrupts adipogenesis and thermogenesis through the activation of Atf3.

**Fig. 4.**
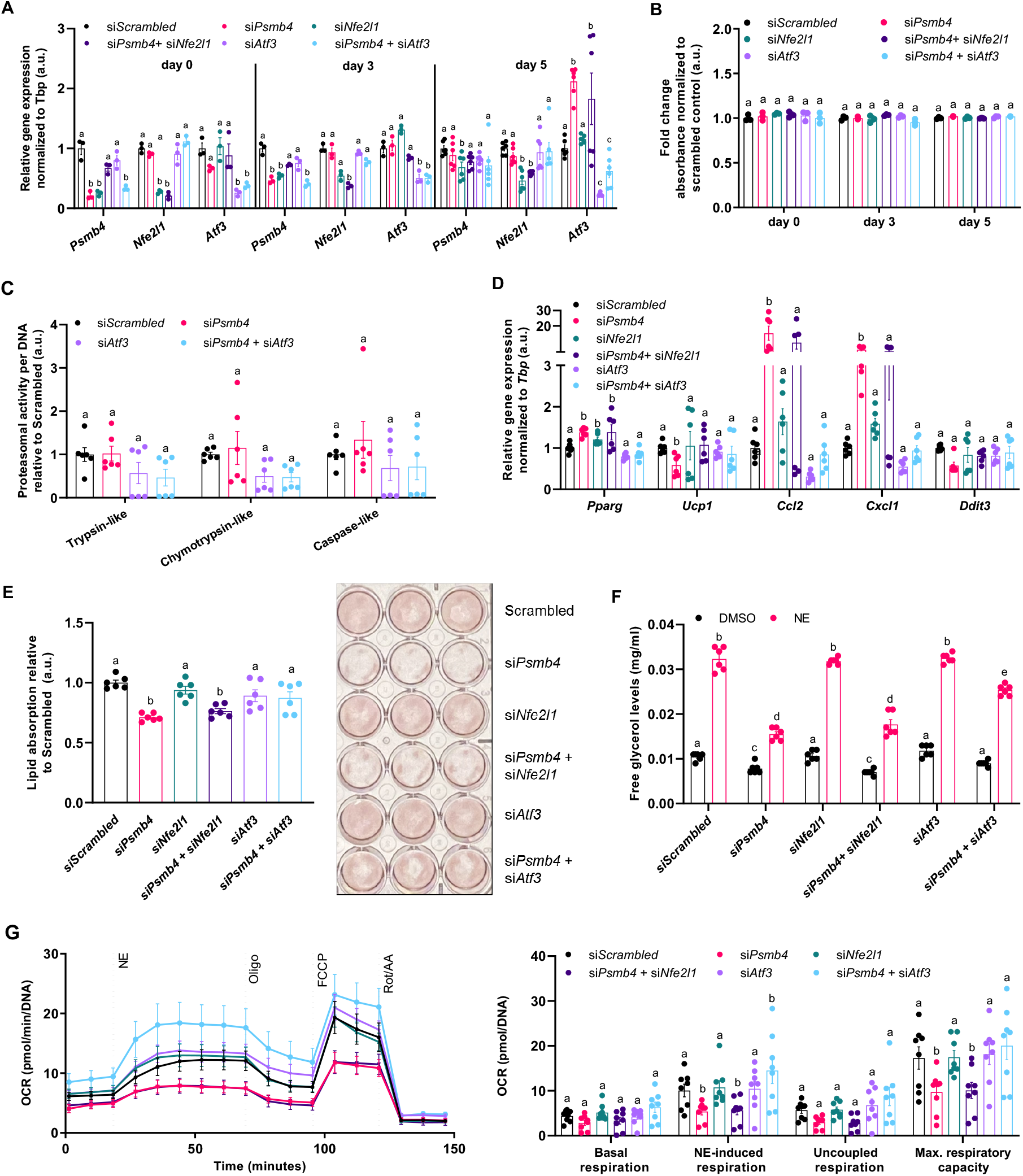
Loss of Psmb4 induces inflammation and blocks adipogenesis via Atf3. **(A)** Viability in adipocytes. **(B)** Relative gene expression of *Psmb4, Nfe2l1*, and *Atf3*. **(C)** Trypsin-like, chymotrypsin-like, and caspase-like proteasome activity in adipocytes (day 5), normalized to DNA content **(D)** Relative gene expression of adipogenesis and stress markers in adipocytes (day 5) after knockdown. **(E)** Oil-Red-O staining in adipocytes after knockdown. **(F)** Supernatant free glycerol levels after treatment with DMSO or 1 μM Norepinephrine (NE) for 1 h. **(G)** Oxygen consumption rate (OCR) in adipocytes after knockdown, normalized to DNA levels (n = 8, from 2 experiments). n = 6 independent measurements from 2 separate experiments. Data are mean ± SEM. Significant if *P* < 0.05, indicated by (*) or different letters.

## 4. Discussion

The proteasome is considered a prerequisite for cellular proteostasis, but its remodeling in response to processes of physiologic adaptation remains largely unexplored. CANDLE/PRAAS syndrome has shed light on the relevance of the proteasome in human pathology, but the underlying molecular mechanisms linking disturbed proteasome function to e.g., inflammation and lipodystrophy remains unclear. Here, we show that in mouse brown adipocytes loss of Psmb4, disturbs proteostasis and protein quality control and, thus, impacts adipogenesis and thermogenesis, whereas Psmb8 silencing had no effect. In these cells, Psmb4 is a constitutively expressed gene that is upregulated during adipogenesis and states of thermogenic activation. Psmb4 is a target gene of Nfe2l1 and this regulation is in line with the general requirement of increased proteasome function during cold adaptation (15). Contrasting this, Psmb8 encodes for a subunit of the immunoproteasome subunit, which is induced by interferons. Psmb8 ablation was previously shown to disrupt proteostasis in immune response (5,19). As the role of the immunoproteasome is especially important in hematopoietic cells (23,24), it is perhaps logical that Psmb8 expression diminishes during differentiation, as there is little requirement for a mature adipocyte. Considering this, it is not surprising that Psmb8 is dispensable for brown adipocyte development and function.

CANDLE/PRAAS patients with mutations in *PSMB4* or *PSMB8* have a similar disease presentation (5) and suffer from lipodystrophy and fevers. Our data suggest that the intrinsic cellular pathways may vary in these patients. It is possible that lipodystrophy in patients with *PSMB8* mutations is a result of autoinflammation and its systemic effects on adipose tissue whereas lipodystrophy in patients with *PSMB4* mutations is caused by a combination of the autoinflammatory syndrome and intrinsically impaired adipogenesis. Our results do not suggest that hyperactivation of BAT is involved in the fever symptoms in patients, as loss of Psmb4 function in mouse brown adipocytes diminishes NST. Nevertheless, altered thermoregulation in the absence of NST might contribute to the symptoms. In addition, we are aware that silencing of genes by RNAi does not completely reflect the natural effects of missense, truncation, or deletion mutations in patients. However, our study warrants further investigation of *PSMB4* mutations on proteasome and adipocyte function. A more complete understanding of systemic as well as cell-specific effects, could improve treatment options for patients.

An interesting and potentially therapeutically relevant aspect of our work is the adaptive activation of Nfe2l1 in response to loss of Psmb4 in adipocytes. This recruitment of Nfe2l1 is most likely a response to insufficient turnover of ubiquitinated proteins caused by proteasome dysfunction, as seen in CANDLE/PRAAS patients (25) or by inhibiting proteasome activity with chemical inhibitors (12). While the activation of Nfe2l1 partly restored total proteasomal activity, we found that this activation of Nfe2l1 was insufficient to overcome the defects in adipogenesis and adipocyte function. Perhaps, while both the loss of Psmb4 and Nfe2l1 cause abnormal proteasomal function, a major difference between the two conditions is the presence of incomplete proteasome intermediates. Reduced total proteasomal activity in the absence of Nfe2l1 was not associated with incomplete proteasome intermediates, and, consequently, also not with a block in adipogenesis. More work is needed to characterize the complex remodeling of cellular proteasome function under proteotoxic stress conditions. Nfe2l1 activation might serve as a therapeutic approach to overcome some of the pathologies associated with PRAAS.

A major finding of our study is that silencing of Atf3 largely rescued the adipocyte defects caused by loss of Psmb4. Atf3 is a member of the mammalian activation transcription factor/cAMP responsive element-binding (CREB) protein family of transcription factors that responds to various stressors (22) and is a downstream target of Atf4 (26). Interestingly, Atf3 was previously shown to downregulate adipogenesis markers and to protect against diet-induced obesity in mice (27). In the context of our research, Atf3 could be viewed as a brake that responds to proteotoxic stress caused by loss of Psmb4. Atf3 does not seem to signal back to the UPS or predispose the cell death, yet its activation induces inflammatory pathways, potentially sending out “danger” and “help” signals. Removing Atf3 clears the path for adipogenesis, lipid metabolism and thermogenesis and might be of therapeutic interest to tackling symptoms associated with PRAAS.

More generally, in the broader context of metabolism, our data underscore the relevance of UPS-mediated protein quality control in maintaining cellular health and function, exemplified here in the context of adipocytes. We show that proteostasis and lipid metabolism are intricately linked in adipocytes and failure to secure proteostasis results in diminished adipogenesis. An important hallmark of obesity-induced adipocyte dysfunction is cellular stress and inflammation, which are tightly linked to aberrant lipid metabolism and insulin resistance. Our finding that maladaptation of UPS and the activation of stress sensors, including Atf3, impede adipogenesis should also be interpreted in the context of potentially hampering obesity-induced adipose tissue expansion or, as in the case of PRAAS, resulting in lipodystrophy. Identifying the key nodes linking proteostasis to cellular stress pathways will represent an important step towards understanding pathological alterations that result in aberrant metabolism, inflammation, premature ageing, and cancer.

## Acknowledgments

The authors thank the members of the Bartelt Lab for their support and for providing an engaging lab environment. The authors thank Brice Emanuelli for providing the WT-1 cell line, Silvia Weidner and Thomas Pitsch for their assistance, and Carolin Muley for critically reading the manuscript. We apologize to colleagues whose work we were not able to cite due to space limitations.

## Author Contributions

N.W. and I.A. performed the experiments and analyzed data. M.S.T. and E.K. provided the mouse model and mouse samples. N.W. and A.B. conceptually designed the study, interpreted the data, and wrote the manuscript. All authors read and commented on the manuscript.

## Figures

**Supplemental fig. S1.**
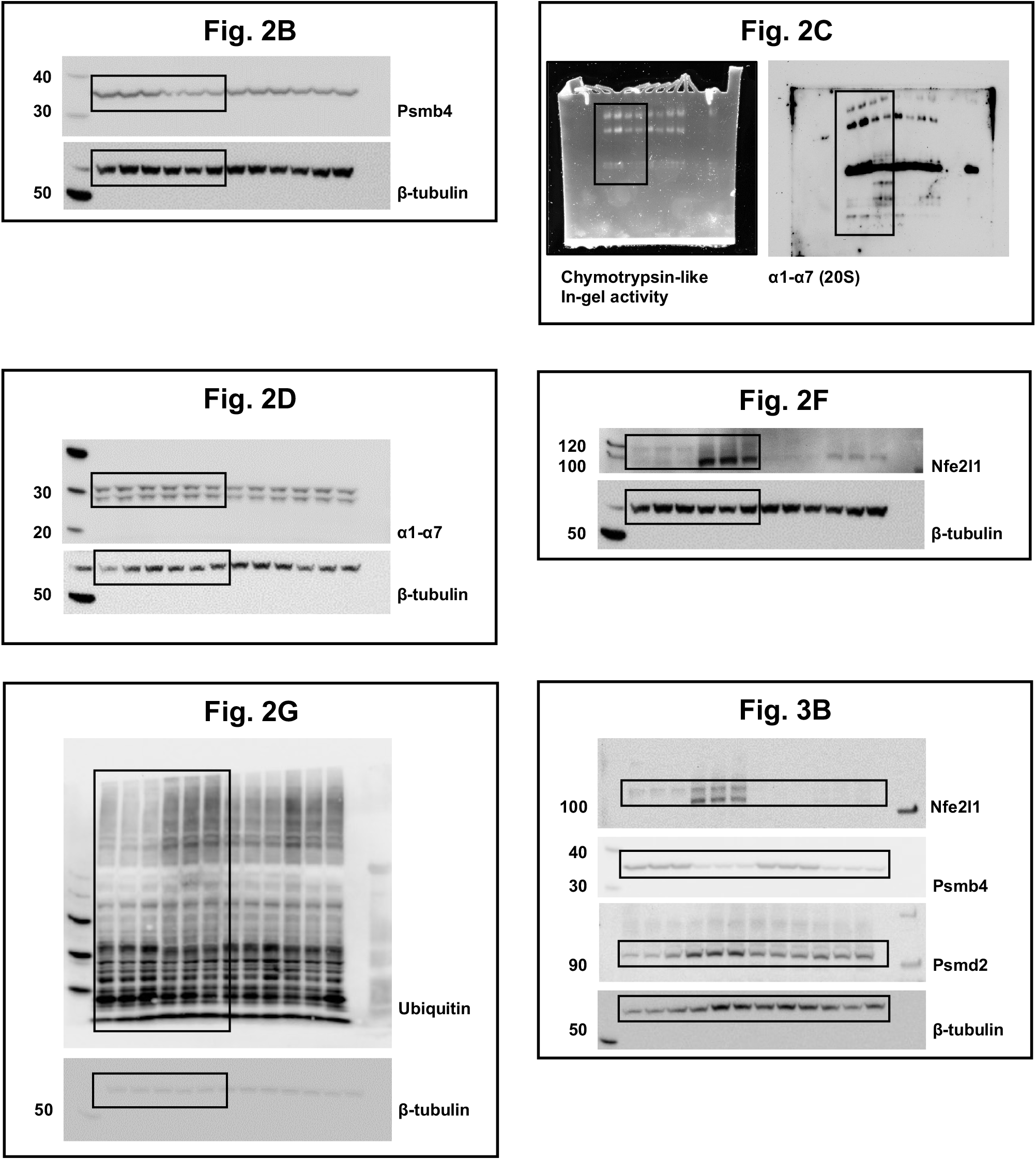
Uncropped immunoblots.

**Supplemental fig. S2.**
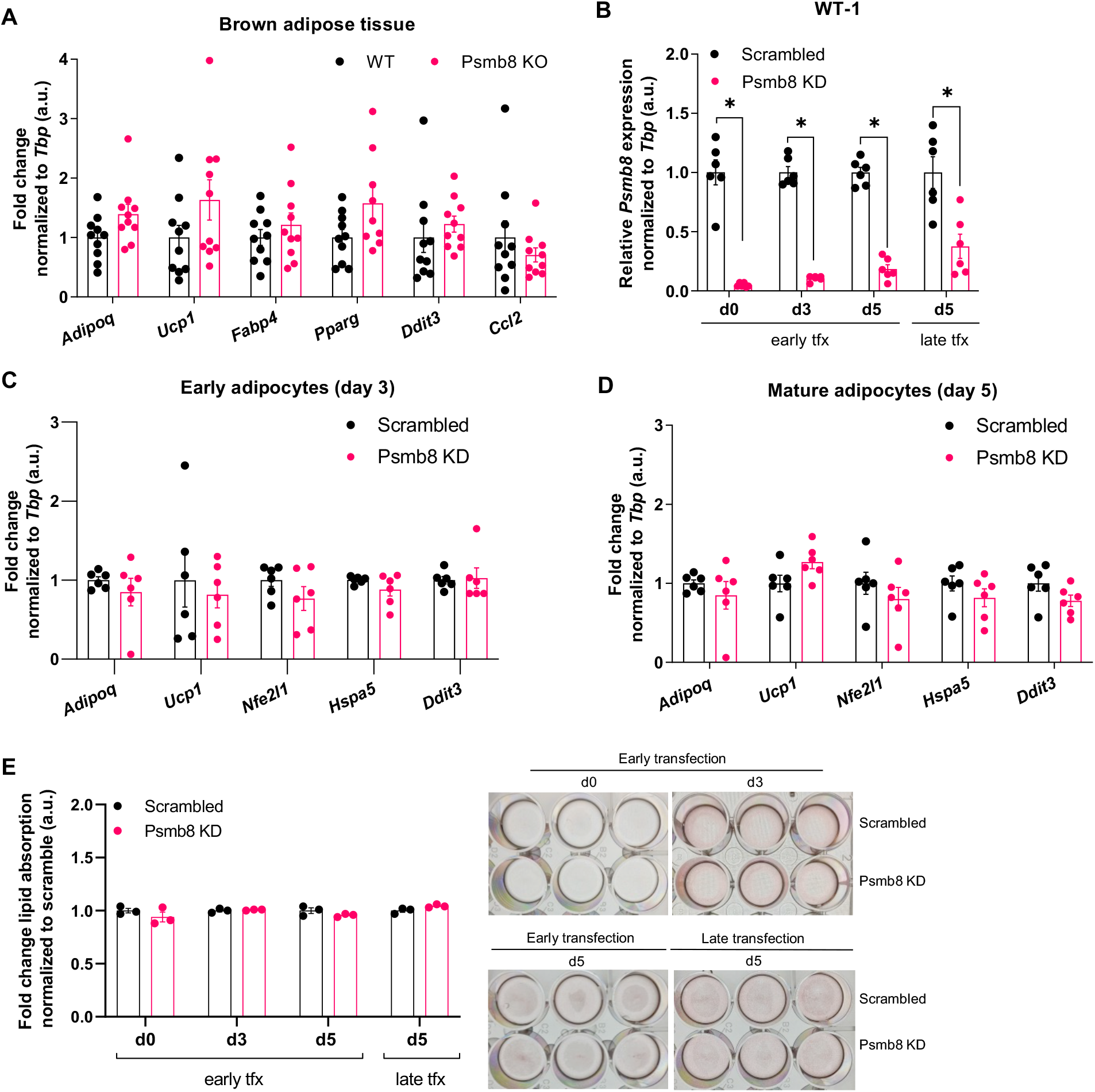
Psmb8 does not affect adipogenesis or cellular function. **(A)** Relative gene expression of adipogenesis and stress markers in BAT from whole-body knock-out mice (n = 11 WT, n = 11 KO mice). **(B)** Relative gene expression of *Psmb8* in mature WT-1 cells after knockdown at different timepoints. (n = 6 from 2 independent experiments). **(C)** Relative gene expression of adipogenesis and stress markers in early WT-1 cells after early knockdown (n = 6 from 2 independent experiments). **(D)** Relative gene expression of adipogenesis and stress markers in mature WT-1 cells after early knockdown (n = 6 from 2 independent experiments). **(E)** Oil Red O staining and lipid resorption in WT-1 cells after knockdown at different timepoints (n=3). Data are mean ± SEM. Significant if *P* < 0.05, indicated by (*) or different letters.

**Supplemental tab. 1.**
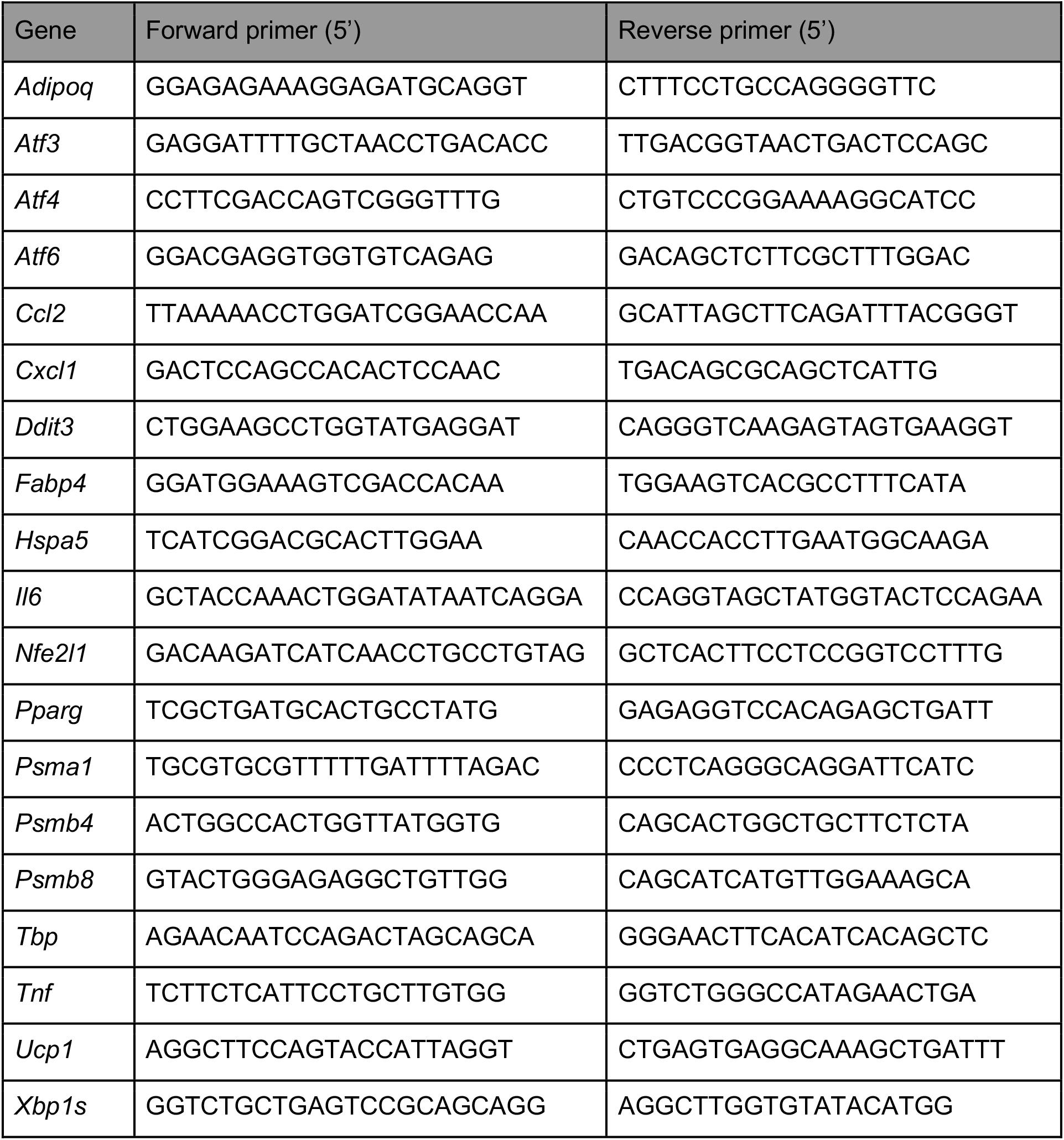
List of primers used for quantitative real-time PCR.

## References

1. Torrelo A, Patel S, Colmenero I, Gurbindo D, Lendínez F, Hernández A, et al. Chronic atypical neutrophilic dermatosis with lipodystrophy and elevated temperature (CANDLE) syndrome. J Am Acad Dermatol. 2010 Mar;62(3):489–95.

2. Agarwal AK, Xing C, DeMartino GN, Mizrachi D, Hernandez MD, Sousa AB, et al. PSMB8 Encoding the β5i Proteasome Subunit Is Mutated in Joint Contractures, Muscle Atrophy, Microcytic Anemia, and Panniculitis-Induced Lipodystrophy Syndrome. Am J Hum Genet. 2010 Dec;87(6):866–72.

3. Kanazawa N. Nakajo-Nishimura Syndrome: An Autoinflammatory Disorder Showing Pernio-Like Rashes and Progressive Partial Lipodystrophy. Allergol Int. 2012;61(2):197–206.

4. Kitamura A, Maekawa Y, Uehara H, Izumi K, Kawachi I, Nishizawa M, et al. A mutation in the immunoproteasome subunit PSMB8 causes autoinflammation and lipodystrophy in humans. J Clin Invest. 2011 Oct 3;121(10):4150–60.

5. Brehm A, Liu Y, Sheikh A, Marrero B, Omoyinmi E, Zhou Q, et al. Additive loss-of-function proteasome subunit mutations in CANDLE/PRAAS patients promote type I IFN production. J Clin Invest. 2015 Oct 20;125(11):4196–211.

6. Ebstein F, Poli Harlowe MC, Studencka-Turski M, Krüger E. Contribution of the Unfolded Protein Response (UPR) to the Pathogenesis of Proteasome-Associated Autoinflammatory Syndromes (PRAAS). Front Immunol. 2019 Nov 26;10:2756.

7. Finley D. Recognition and Processing of Ubiquitin-Protein Conjugates by the Proteasome. Annu Rev Biochem. 2009 Jun;78(1):477–513.

8. Collins GA, Goldberg AL. The Logic of the 26S Proteasome. Cell. 2017 May;169(5):792– 806.

9. Kimura H, Caturegli P, Takahashi M, Suzuki K. New Insights into the Function of the Immunoproteasome in Immune and Nonimmune Cells. J Immunol Res. 2015;2015:1–8.

10. Huber EM, Basler M, Schwab R, Heinemeyer W, Kirk CJ, Groettrup M, et al. Immuno- and Constitutive Proteasome Crystal Structures Reveal Differences in Substrate and Inhibitor Specificity. Cell. 2012 Feb;148(4):727–38.

11. Mann JP, Savage DB. What lipodystrophies teach us about the metabolic syndrome. J Clin Invest. 2019 Aug 5;129(10):4009–21.

12. Cannon B, Nedergaard J. Brown Adipose Tissue: Function and Physiological Significance. Physiol Rev. 2004 Jan;84(1):277–359.

13. Bartelt A, Bruns OT, Reimer R, Hohenberg H, Ittrich H, Peldschus K, et al. Brown adipose tissue activity controls triglyceride clearance. Nat Med. 2011 Feb;17(2):200–5.

14. Lemmer IL, Willemsen N, Hilal N, Bartelt A. A guide to understanding endoplasmic reticulum stress in metabolic disorders. Mol Metab. 2021 Jan;101169.

15. Bartelt A, Widenmaier SB, Schlein C, Johann K, Goncalves RLS, Eguchi K, et al. Brown adipose tissue thermogenic adaptation requires Nrf1-mediated proteasomal activity. Nat Med. 2018 Mar;24(3):292–303.

16. Radhakrishnan SK, Lee CS, Young P, Beskow A, Chan JY, Deshaies RJ. Transcription Factor Nrf1 Mediates the Proteasome Recovery Pathway after Proteasome Inhibition in Mammalian Cells. Mol Cell. 2010 Apr;38(1):17–28.

17. Sha Z, Goldberg AL. Proteasome-Mediated Processing of Nrf1 Is Essential for Coordinate Induction of All Proteasome Subunits and p97. Curr Biol. 2014 Jul;24(14):1573–83.

18. Fehling HJ, Swat W, Laplace C, Kühn R, Rajewsky K, Müller U, et al. MHC Class I Expression in Mice Lacking the Proteasome Subunit LMP-7. Science. 1994 Aug 26;265(5176):1234–7.

19. Seifert U, Bialy LP, Ebstein F, Bech-Otschir D, Voigt A, Schröter F, et al. Immunoproteasomes Preserve Protein Homeostasis upon Interferon-Induced Oxidative Stress. Cell. 2010 Aug;142(4):613–24.

20. Yazgili AS, Meul T, Welk V, Semren N, Kammerl IE, Meiners S. In-gel proteasome assay to determine the activity, amount, and composition of proteasome complexes from mammalian cells or tissues. STAR Protoc. 2021 Jun;2(2):100526.

21. Rull A, Camps J, Alonso-Villaverde C, Joven J. Insulin Resistance, Inflammation, and Obesity: Role of Monocyte Chemoattractant Protein-1 (orCCL2) in the Regulation of Metabolism. Mediators Inflamm. 2010;2010:1–11.

22. Ku H-C, Cheng C-F. Master Regulator Activating Transcription Factor 3 (ATF3) in Metabolic Homeostasis and Cancer. Front Endocrinol. 2020 Aug 14;11:556.

23. Çetin G, Klafack S, Studencka-Turski M, Krüger E, Ebstein F. The Ubiquitin–Proteasome System in Immune Cells. Biomolecules. 2021 Jan 5;11(1):60.

24. Tubío-Santamaría N, Ebstein F, Heidel FH, Krüger E. Immunoproteasome Function in Normal and Malignant Hematopoiesis. Cells. 2021 Jun 22;10(7):1577.

25. Sotzny F, Schormann E, Kühlewindt I, Koch A, Brehm A, Goldbach-Mansky R, et al. TCF11/Nrf1-Mediated Induction of Proteasome Expression Prevents Cytotoxicity by Rotenone. Antioxid Redox Signal. 2016 Dec;25(16):870–85.

26. Forsström S, Jackson CB, Carroll CJ, Kuronen M, Pirinen E, Pradhan S, et al. Fibroblast Growth Factor 21 Drives Dynamics of Local and Systemic Stress Responses in Mitochondrial Myopathy with mtDNA Deletions. Cell Metab. 2019 Dec;30(6):1040-1054.e7.

27. Ku H-C, Chan T-Y, Chung J-F, Kao Y-H, Cheng C-F. The ATF3 inducer protects against diet-induced obesity via suppressing adipocyte adipogenesis and promoting lipolysis and browning. Biomed Pharmacother Biomedecine Pharmacother. 2022 Jan;145:112440.

